# High-Dimensional Spectral Cytometry Reveals Therapeutically Relevant Immune Subtypes in Gastric Cancer

**DOI:** 10.1101/2023.03.29.534765

**Authors:** Miseker Abate, Teng Fei, Ya Hui Lin, Shoji Shimada, Harrison Drebin, Eunise Chen, Laura Tang, Vivian E Strong, Santosha A. Vardhana

## Abstract

Identification of locally advanced gastric cancer (GC) patients who might potentially benefit from immune-based strategies is limited by both the poor predictive quality of existing biomarkers, including molecular subtypes, tumor mutational burden, and PD-L1 expression, as well as inadequate understanding of the gastric cancer immune microenvironment. Here, we leveraged high-dimensional spectral cytometry to re-classify locally advanced gastric tumors based on immune composition. The gastric cancer microenvironment was comprised of a diverse immune infiltrate including high proportions of plasmablasts, macrophages, and myeloid-derived suppressor cells. Computational cell typing and sample clustering based on tiered broad immune and T-cell focused phenotyping identified three distinct immune subtypes. The most immunogenic subtype exhibited high proportions of activated CD4+ T-cells and plasmablasts and included tumors that would have been classified as non-immunogenic based on prior classifications. Analysis of gastric cancer patients treated with immune checkpoint blockade indicates that patients who responded to immunotherapy had a pre-treatment tumor composition that corresponded to higher immune scores from our analysis. This work establishes a novel immunological classification of gastric cancer including identification of patients and immune networks likely to benefit from immune-based therapies.

## Introduction

Gastric cancer (GC) is the third leading cause of cancer-related deaths worldwide with over 1 million cases diagnosed annually in the United States.^1^ GC is a deadly disease with a 5-year overall survival rate of less than 40% for patients with locally advanced disease and less than 20% for patients with advanced stage disease.^2^ Immunotherapy is a promising therapeutic approach in GC with disease specific and overall survival benefits seen in the ATTRACTION-2^3^, CheckMate-649^4^ and KEYNOTE-811^5^ clinical trials. However, most GC tumors remain insensitive to immunotherapy and there is currently no classification system or biomarker that can consistently identify patients who respond to immune based therapies.

To date, the only clinically applicable biomarkers and classification systems employed with respect to immunogenicity in GC are elevated PD-L1 scores and the Cancer Genome Atlas (TCGA) group’s comprehensive molecular categorization of GC.^6^ The TCGA noted that Epstein Barr virus (EBV) and microsatellite instability high (MSI-high) GC were more immunogenic compared with the more common chromosomal instability (CIN) and genomically stable (GS, corresponding to diffuse histology) subtypes based on their expression of *CD274* and *PDCD1LG2*, which encode the immunotherapy targets PD-L1 and PD-L2.^6^ However, the survival benefit of pembrolizumab in the KEYNOTE-062 trial was similar across pre-specified GC subgroups including diffuse type,^7^ and the KEYNOTE-811 study did not show a significant difference in response rate between patients with positive and negative PD-L1 scores.^5^

To overcome these limitations, we leveraged high-dimensional spectral cytometry to categorize gastric cancer based on intra-tumoral immune signatures. Our results establish novel immune signatures in gastric cancer and identify intercellular immune correlations that may serve as biomarkers for immunogenicity.

## Results

### Patient characteristics and demographics

Fresh gastric tumor (n=38), adjacent normal (n=29), and lymph node (n=26) samples were obtained from 38 patients who underwent resection of gastric tumors or esophagogastric duodenoscopy (EGD) with biopsy at Memorial Sloan Kettering Cancer Center (MSK) between 2021-2022 (**Fig, 1A, clinicopathologic characteristics are included in Supplementary Table 1**). Most patients were male (26/38) and of self-identified white race (26/38). Tumors were most frequently of the intestinal (15/38) and mixed (13/38) pathologic subtypes. Most tumors were of later T stages (27/38) and of N1 or greater nodal stage (15/28). Tumors had a wide distribution of PD-L1 scores.

### Distinct immune intra-tumoral landscape of gastric cancers

We first performed broad immune phenotyping of CD45+ immune cells in gastric tumors, adjacent normal mucosa, and draining lymph nodes. As control, five melanoma tumors were profiled using identical methodology (**Fig. 1A**). Gastric tumors exhibited a rich immune infiltration, with an increased proportion of CD45+ cells compared to adjacent normal tissue and at a similar density to melanomas (**Supplementary Fig. 1A**). This increased infiltrate was driven mainly by myeloid cells, as gastric tumors exhibited significantly higher proportions of dendritic cells (DCs), macrophages, and myeloid-derived suppressor cells (MDSC) compared to adjacent normal tissue but similar proportions of T, B and NK-cells (**Fig. 1B-C** **and Supplementary Fig. 1C**). This immune composition was also distinct from melanomas, which demonstrated a similar proportion of T, B, and NK cells but more DCs and fewer macrophages or MDSCs (**Fig. 1B-C and Supplementary Fig. 1C**).

**Figure 1.**
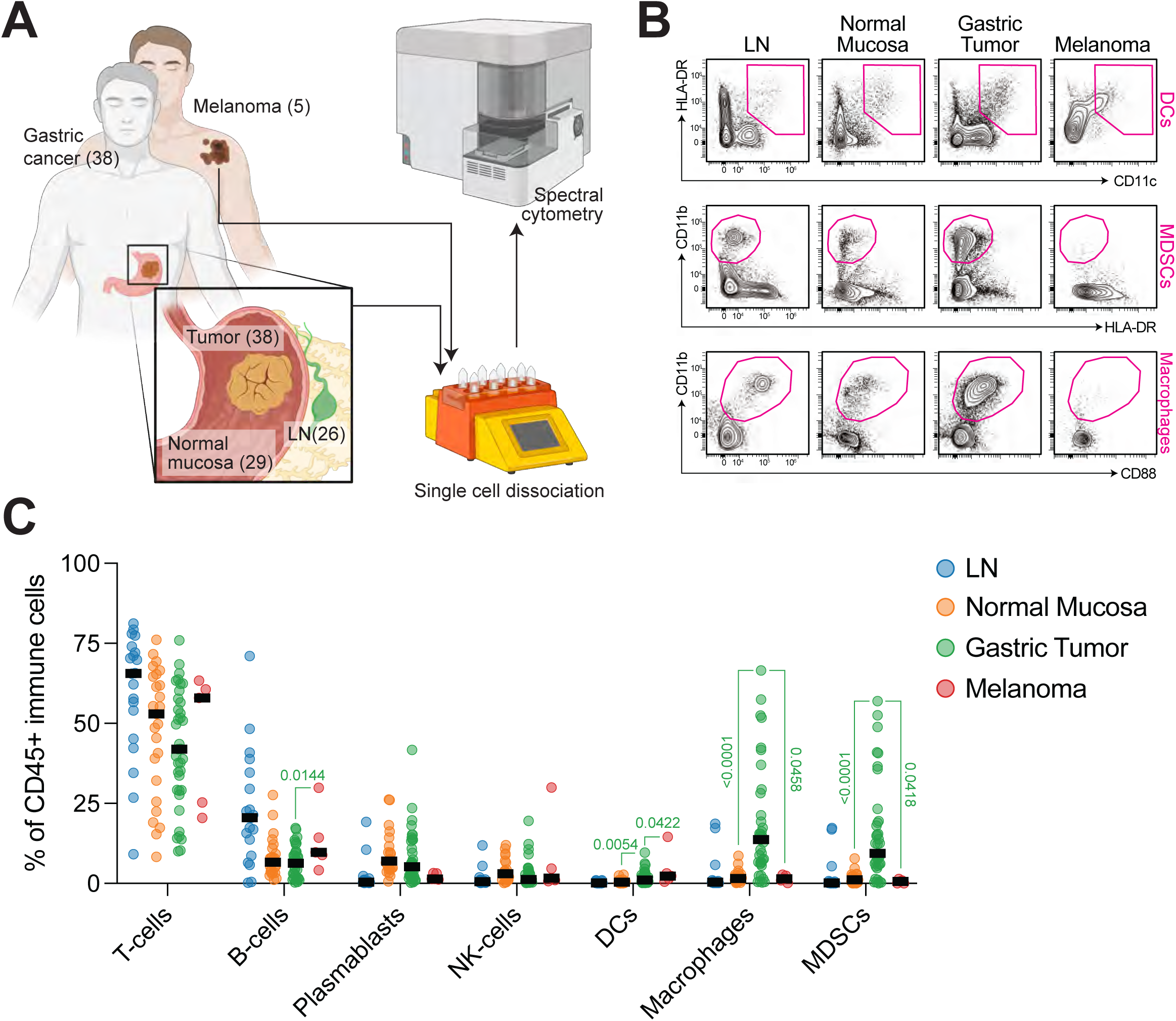
Distinct tumor-specific landscape of localized gastric adenocarcinoma. (a) Sample acquisition strategy for gastric cancer and melanoma patients. (b) Contour plots of myeloid and DC cell abundances in gastric tumors, normal mucosa, and lymph nodes compared with melanoma tumors. (c) Quantification of relative abundance of CD45+ cell subsets in gastric tumors, normal mucosa, lymph nodes and melanoma tumors. P values as indicated for gastric tumors were calculated by one-way ANOVA with Sidak’s multiple comparisons post-test relative to normal mucosa and by student’s t-test relative to melanoma tumors.

### Gastric tumors are marked by altered T-cell differentiation

We next interrogated the T-cell landscape of gastric tumors using a parallel high-dimensional flow cytometry panel focused on identifying CD4+ and CD8+ T-cell subsets (**Supplementary Fig. 2A**). As expected, CD8+ T-cells comprised a higher proportion of CD45+ immune cells within gastric tumors as compared to draining lymph nodes, resulting in a decreased CD4:CD8 ratio within tumors (**Supplementary Fig. 2B**). Intra-tumoral CD8+ T-cells were both more activated and more proliferative than adjacent normal tissue but notably decreased compared to melanomas, suggesting a less potent CD8+ T-cell response in gastric tumors (**Fig. 2A-B**). Interestingly, intra-tumoral CD8+ T-cells in melanoma showed hallmarks of T-cell exhaustion, with significant induction of the exhaustion-associated transcription factor TOX and expression of inhibitory receptors including PD-1 and CD39 (**Fig. 2A**).^8–11^ Conversely, intratumoral CD8+ T-cells in gastric cancer did not significantly induce TOX expression and had limited expression of immune checkpoints, suggesting that CD8+ T-cell responses might be suppressed in gastric cancer via mechanisms other than PD-1/PD-L1 interactions.^12, 13^

**Figure 2.**
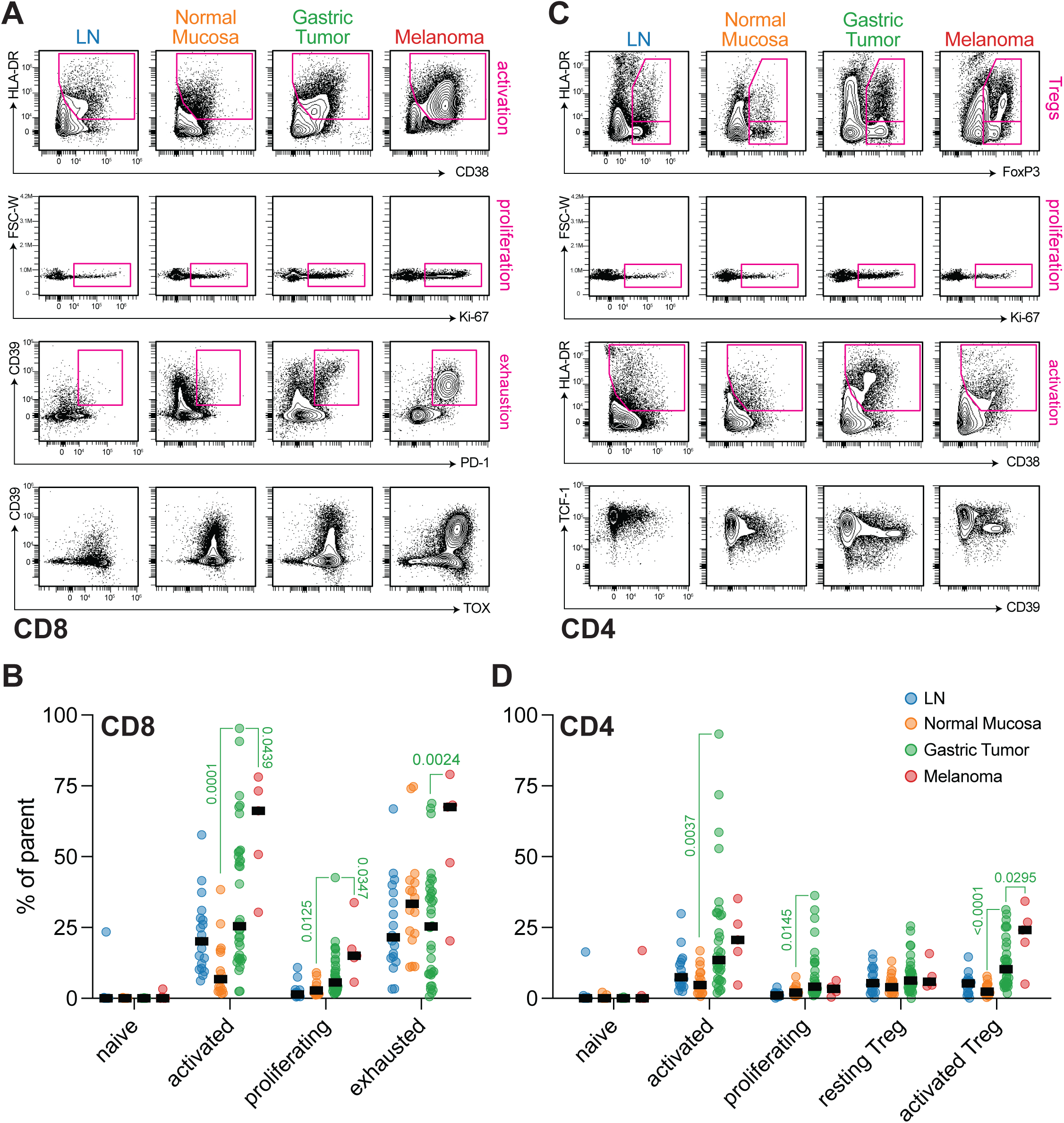
Robust CD4+ and CD8+ T-cell activation in gastric adenocarcinoma. (a,b) Contour plots (a) and quantification (b) of CD8+ T-cell states in gastric tumors, normal mucosa, lymph nodes and melanomas. (c,d) Contour plots (c) and quantification (d) of CD4+ T-cell states in gastric tumors, normal mucosa, lymph nodes and melanomas. P values as indicated for gastric tumors were calculated by one-way ANOVA with Sidak’s multiple comparisons post-test relative to normal mucosa and by student’s t-test relative to melanoma tumors.

Gastric tumors did, however, show signs of a potent anti-tumor CD4+ T-cell response. Gastric tumors exhibited a robust increase in activated and proliferating CD4+ T-cells that was similar to that seen in melanoma tumors (**Fig. 2C-D**). In CD4+ T-cells, downregulation of TCF-1 is associated with effector differentiation,^14^ and we observed the highest frequency of TCF-1 loss in gastric tumor-associated CD4+ T-cells (**Fig. 2C**). Finally, while we did observe tumor-specific accumulation of activated regulatory T-cells (Tregs) in gastric cancers, significantly fewer activated Tregs were seen as compared to melanomas (**Fig. 2C-D**). Taken together, these results suggest that CD4+ T-cells play a significant role in gastric cancer immune surveillance.

### Unbiased clustering pipeline identifies distinct immune archetypes in gastric tumors

Given the unique immune characteristics of gastric tumors and the inability of traditional metrics to predict response to immunotherapy in gastric cancer, we next took an unbiased approach to classify gastric tumors based on immune landscape alone. Cell types were determined by mini-batch K-means clustering of mean fluorescence intensity for each fluorophore, where the number of distinct cell clusters were selected by the Elbow method on the within-cluster sum of square curve.^15, 16^ Through this method 14 distinct cell clusters were identified in the broad immune panel and 13 cell clusters in the T-cell panel (**Fig. 3A-B and Supplementary Fig. 3A-B**). Hierarchical clustering of the derived cell composition for the broad immune panel alone, T-cell panel alone, or combined data from both panels identified three immune subtypes, with tiered immune cell composition indicating least immunogenic (IS1), immunogenic (IS2), and most immunogenic groups (IS3) (**Fig. 3C and supplementary Fig. 3C**). Clustering based on either broad immune or T-cell phenotyping identified four groups, with similar graded distribution of the joint immune clusters across broad immune and T cell panel-based sub-groups (**Supplementary Fig. 3D**).

**Figure 3.**
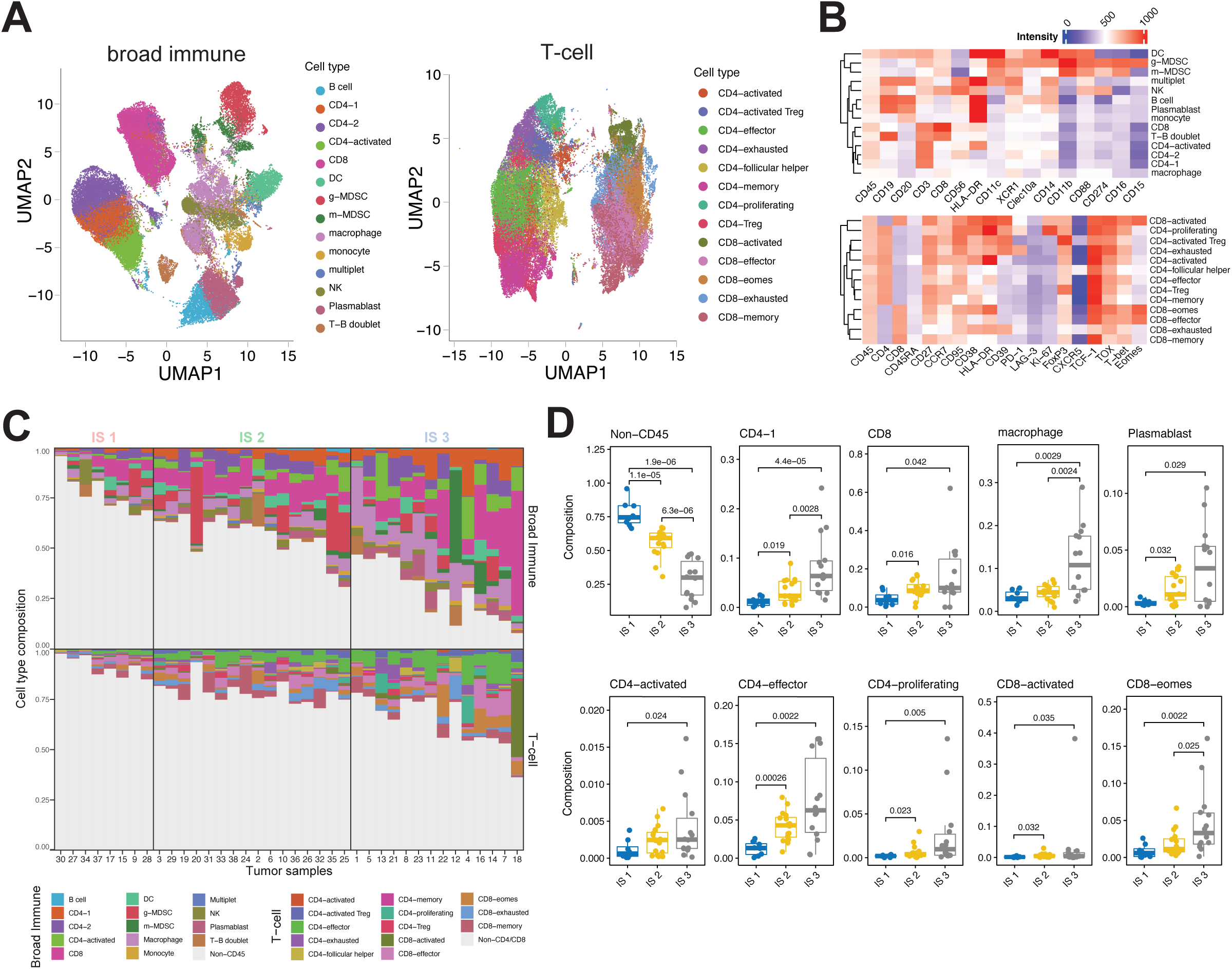
Hierarchical clustering identifies novel tumor-specific immune subtypes in gastric adenocarcinoma. (a) UMAP projection of CD45+ immune cells based on broad immune panel (left) and CD4+ and CD8+ lymphocytes based on T-cell panel (right). Cell type annotations as indicated. (b) Heatmap displaying relative fluorescence intensity of fluorophores within each mini-batch K-means derived cell type as indicated. (c) Distribution of patient-specific cell type frequency stratified by immune subtypes obtained by hierarchical clustering. (d) Frequency of cell type by immune score as a proportion of viable cells. P values as indicated were calculated by Wilcoxon’s rank-sum test.

Further characterization of the cell compositions in the immune subtypes (IS) revealed discrete cell types enriched in immunogenic tumors (**Fig. 3D**). As expected, immunogenic tumors had increasing proportions of CD45+ immune cells. Broadly, immunogenic tumors were associated with higher proportions of CD4+ and CD8+ T-cells as well as macrophages and plasma blasts, but not dendritic cells or NK cells (**Fig. 3D and Supplementary Fig. 3E**). Within the T-cell panel, IS2 and IS3 exhibited high rates of effector and proliferating CD4+ T-cells and activated T-regs as well as activated and Eomes-high CD8+ T-cells but not higher rates of effector or exhausted CD8+ T-cells (**Fig. 3D and Supplementary Fig.3F**). Notably, tumor-specific immune subtypes were not associated with a corresponding infiltrate in tumor-adjacent normal mucosa or draining lymph nodes (**Supplementary Fig. 3G-H**) suggesting that the immune landscape of gastric cancer is tumor-specific.

### Gastric cancer immune subtypes are tumor-specific and independent of somatic mutational profile or PD-L1 status

An immunogenic landscape has been associated with several tumor-specific factors in other cancer subtypes, including tumors with microsatellite instability, high somatic mutational burden, high expression of PD-L1, and/or evidence of oncogenic viral infection.^17–20^ In contrast, we found that unbiased gastric cancer immune subtypes were largely independent of these factors. Apart from EBV-associated tumors, of which only 1 tumor was profiled, all other Lauren subtypes of gastric cancer were observed across immune subtypes (**Fig. 4A-B**). Similarly, we did not observe any significant differences in immune score by TNM stage, except for T4 tumors, which were absent from the least immunogenic subtype (IS1) and M1 tumors, which were present only in the most immunogenic subtype (**Fig. 4C-D and Supplementary Fig. 4A**). Finally, immune subtypes were not dictated by either receipt of prior neoadjuvant chemotherapy or response to neoadjuvant chemotherapy, which was similar in patients across immune subtypes (**Fig. 4E and Supplementary Fig. 4B**).

**Figure 4.**
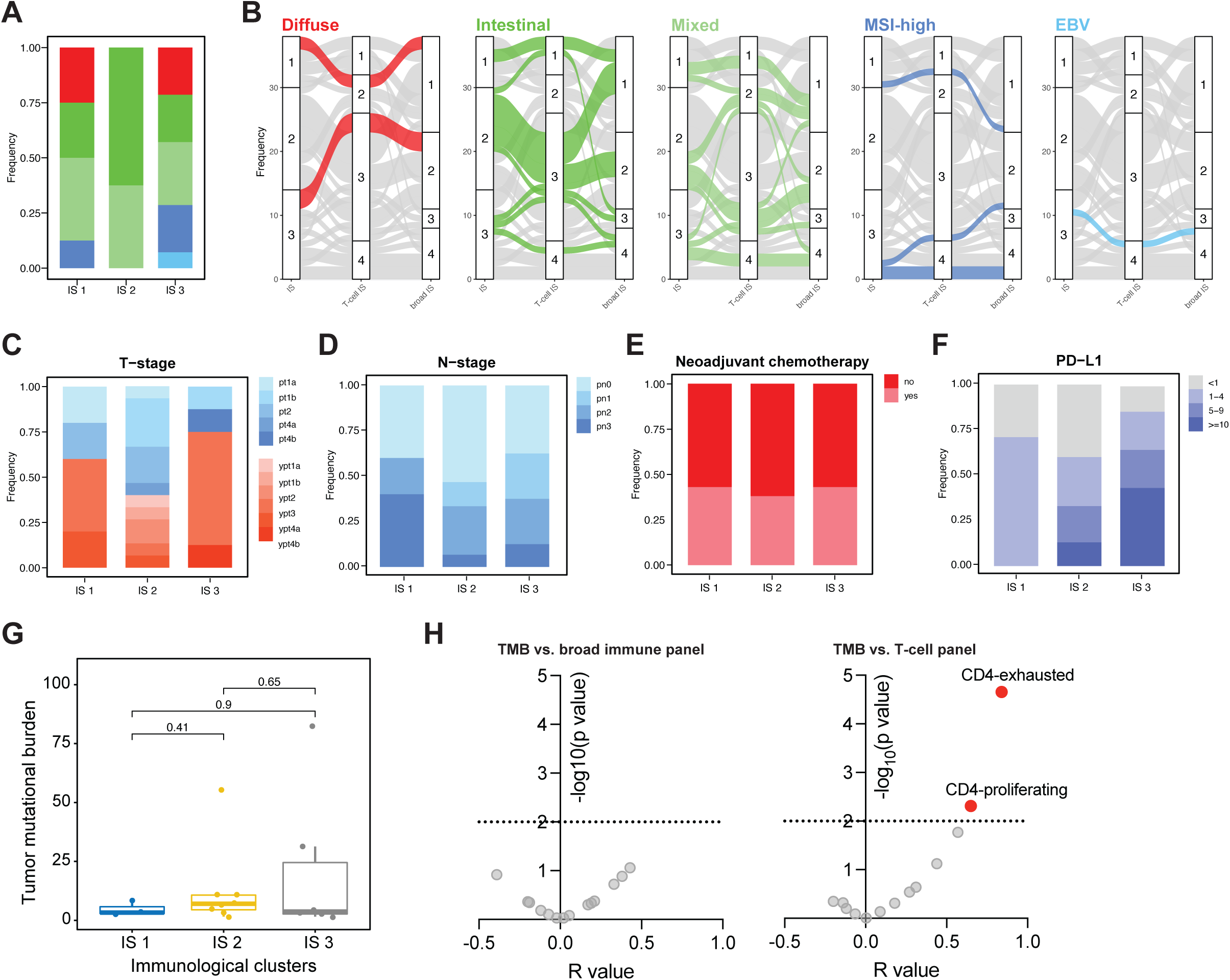
Gastric cancer immune subtypes are independent of other clinical and molecular disease markers. (a) Distribution of gastric cancer molecular and histological subtypes across immune subtypes. (b) Alluvial plots of gastric cancer subtype distribution based on overall immune subtypes, T-cell specific subtypes and broad immune-specific subtypes. (c-f) Distribution of pathologic T stage (c), N stage (d), receipt of Neoadjuvant Chemotherapy (e), and PD-L1 score by IHC (f) across immune subtypes. (g) Tumor mutational burden (TMB) in patients with MSK-IMPACT testing stratified by immune subtype. P values calculated by Wilcoxon’s rank-sum test. (h) Correlation of TMB with frequency of tumor-specific cell types as determined by the broad immune panel (left) or T-cell specific panel (right), plotted by both strength of correlation (R) and p-value for non-zero slope.

Finally, assessment of PD-L1 scores and immune subtypes (**Table 1**) showed the most immunogenic subtype (IS3) to have higher proportions of tumors with PD-L1 scores between 5-9 and > 10 (**Fig. 4F**); however, tumors with PD-L1 scores less than 5 were identified in all three immune subtypes, indicating that PD-L1 score inaccurately predicts immunogenicity in gastric cancer. We did not observe a significant association between tumor mutational burden and immune subtypes (**Fig. 4G**); however, correlation analyses demonstrated that proliferating and exhausted CD4+ T-cells correlated most strongly with tumor mutational burden, suggesting once again that CD4+ T-cells may play a particularly significant role in gastric cancer immune surveillance (**Fig. 4H**).

**Table 1.** Patient characteristics and demographics across immune subtypes based on integrated immune phenotyping

| Characteristic | IS 1, N = 8<sup>1</sup> | IS 2, N = 16<sup>1</sup> | IS 3, N = 14<sup>1</sup> | p-value<sup>2</sup> |
| --- | --- | --- | --- | --- |
| **Age** | 69 (62, 79) | 72 (68, 76) | 68 (60, 75) | 0.6 |
| **BMI\*** | 23.6 (20.6, 26.8) | 26.9 (25.4, 27.6) | 25.5 (20.8, 27.3) | 0.3 |
| **Sex** |  |  |  | 0.8 |
| Female | 2 (25%) | 6 (38%) | 4 (29%) |  |
| Male | 6 (75%) | 10 (62%) | 10 (71%) |  |
| **Race** |  |  |  | 0.9 |
| Asian | 0 (0%) | 3 (21%) | 2 (14%) |  |
| Black | 1 (14%) | 1 (7.1%) | 2 (14%) |  |
| White | 6 (86%) | 10 (71%) | 10 (71%) |  |
| Other* | 1 (12%) | 2 (12%) | 0 (0%) |  |
| **Comorbidities** |  |  |  |  |
| H. Pylori | 0 (0%) | 2 (12%) | 1 (7.1%) | 0.8 |
| GERD* | 2 (25%) | 9 (56%) | 6 (43%) | 0.4 |
| Diabetes Mellitus | 0 (0%) | 1 (6.2%) | 2 (14%) | 0.6 |
| CAD* | 1 (12%) | 3 (19%) | 3 (21%) | >0.9 |
| Smoking history | 1 (12%) | 0 (0%) | 2 (14%) | 0.3 |
| **Neoadjuvant Chemotherapy** | 3 (43%) | 6 (38%) | 6 (43%) | >0.9 |
| **Histologic Grade** |  |  |  | 0.4 |
| Well | 1 (12%) | 1 (6.2%) | 0 (0%) |  |
| Moderate | 2 (25%) | 6 (38%) | 3 (21%) |  |
| Moderate to Poor | 0 (0%) | 3 (19%) | 1 (7.1%) |  |
| Poor | 5 (62%) | 6 (38%) | 10 (71%) |  |
| **Tumor Subtype** |  |  |  | 0.057 |
| Diffuse | 2 (25%) | 0 (0%) | 3 (21%) |  |
| EBV | 0 (0%) | 0 (0%) | 1 (7.1%) |  |
| Intestinal | 2 (25%) | 10 (62%) | 3 (21%) |  |
| Mixed | 3 (38%) | 6 (38%) | 4 (29%) |  |
| MSI-High | 1 (12%) | 0 (0%) | 3 (21%) |  |
| **Pathologic T Stage** |  |  |  | **0.013** |
| pT1a | 1 (20%) | 2 (13%) | 0 (0%) |  |
| pT1b | 0 (0%) | 5 (33%) | 1 (12%) |  |
| pT2 | 1 (20%) | 5 (33%) | 0 (0%) |  |
| pT3 | 2 (40%) | 1 (6.7%) | 5 (62%) |  |
| pT4a | 1 (20%) | 2 (13%) | 0 (0%) |  |
| pT4b | 0 (0%) | 0 (0%) | 2 (25%) |  |
| Unknown | 3 | 1 | 6 |  |
| **Pathologic N Stage** |  |  |  | 0.9 |
| N0 | 2 (40%) | 8 (53%) | 3 (38%) |  |
| >N1 | 3 (60%) | 7 (47%) | 5 (62%) |  |
| Unknown | 3 | 1 | 6 |  |
| **Pathologic M Stage** |  |  |  | **0.048** |
| M0 | 2 (100%) | 5 (100%) | 1 (25%) |  |
| M1 | 0 (0%) | 0 (0%) | 3 (75%) |  |
| Unknown | 6 | 11 | 10 |  |
| **PD-L1** |  |  |  | **0.10** |
| <1 | 2 (29%) | 6 (40%) | 2 (14%) |  |
| 1-4 | 5 (71%) | 4 (27%) | 3 (21%) |  |
| 5-9 | 0 (0%) | 3 (20%) | 3 (21%) |  |
| >=10 | 0 (0%) | 2 (13%) | 6 (43%) |  |
| Unknown | 1 | 1 | 0 |  |

<sup>1</sup> Median (IQR); n (%)<sup>2</sup> Kruskal-Wallis rank sum test; Fisher's exact test
\*BMI: Body Mass Index; *H. Pylori*: *Helicobacter Pylori*; GERD: Gastroesophageal Reflux Disease; Cardiovascular Disease; ASA: American Society of Anesthesiology NAC: Neoadjuvant Chemotherapy; MSI-High: Microsatellite Instability High

### Gastric tumors are characterized by novel cellular networks that predict response to immune checkpoint blockade

Given the unique profile of tumor-infiltrating T-cells in gastric cancer, we next asked whether intratumoral accumulation of specific immune cell types might be associated with increased T-cell infiltration. A correlation analysis of broad immunophenotyping data revealed a highly significant positive correlation between CD8 T-cell and plasmablast infiltration (r=0.80, Bonferonni, p<0.001) which was not significant in either lymph nodes or tumor adjacent normal mucosa (r=0.69 and r=0.50, respectively) (**Fig. 5A and Supplementary Fig. 5A**). This, in combination with our observation that plasmablasts were significantly more enriched in more immunogenic gastric cancer subtypes (**Fig. 3D**), suggested that plasmablasts might play a significant role in recruiting and/or sustaining intratumoral CD8+ T-cell responses in gastric cancer. To validate this finding, we analyzed a recently published dataset comprised of annotated single-cell RNA sequencing data from 48 GC and matched normal samples.^21^ Correlation analysis demonstrated a high correlation of multiple plasma cell subsets with T-cell clusters, with a particularly significant association of IgA-producing plasma cells with T-cell subsets that have been implicated in the response to immune checkpoint blockade, including ‘immune-homing’ CXCL13+ CD8+ (T27) and CD4+ (T06) T-cells and both effector and proliferating CD8+ and CD4+ T-cells (T8, T15, T27) (**Fig. 5B**).^22–25^ This association with IgA-expressing plasma cells is notable given that they are present within normal gastric mucosa, suggesting that immune evasion during gastric carcinogenesis may involve depletion of tissue-resident IgA-producing plasma cells.^26–28^

**Figure 5.**
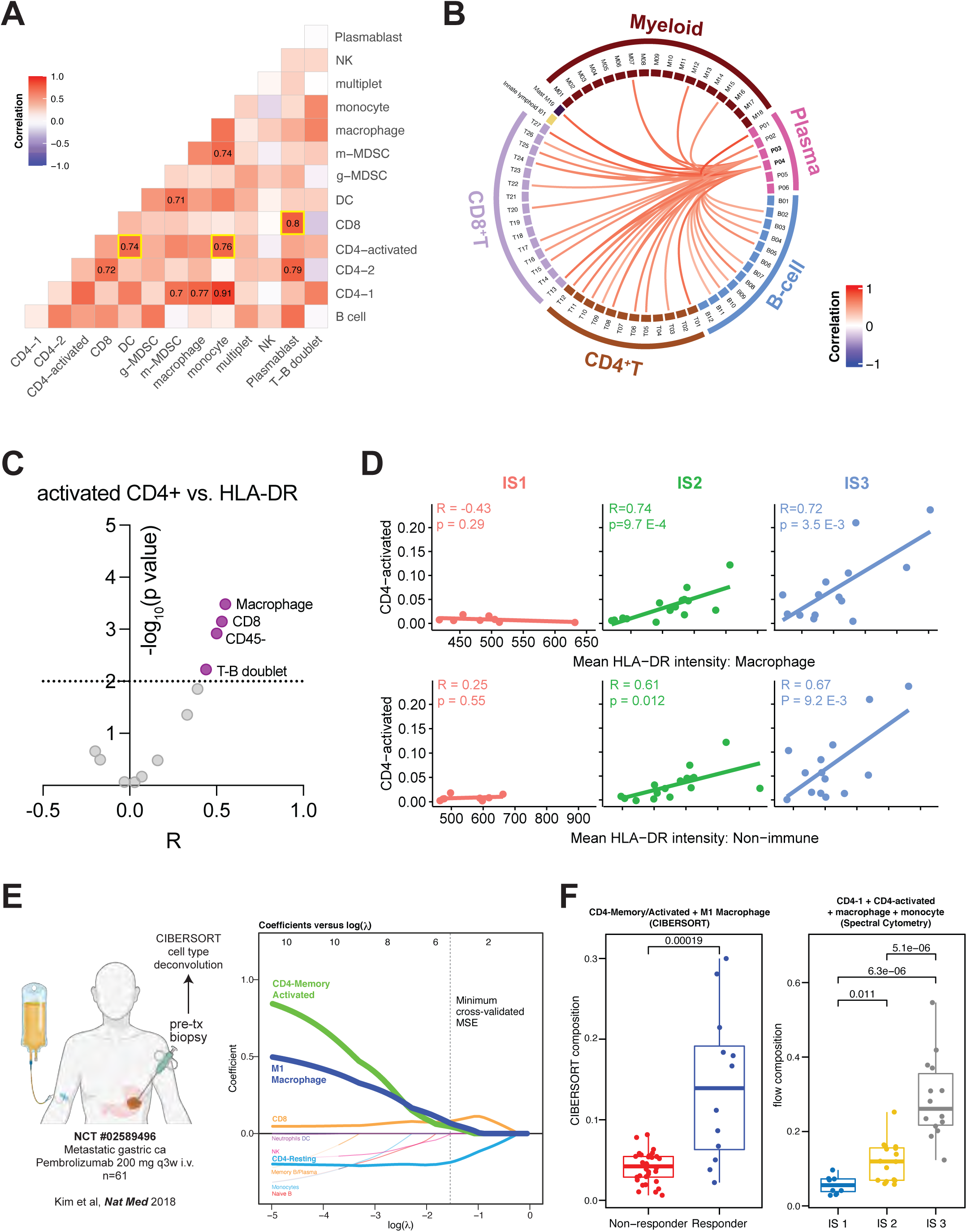
Gastric cancer is characterized by novel immune networks that predict response to immune checkpoint blockade. (a) Correlation analysis of gastric tumor-specific immune cell types. Correlation values significantly different from zero (Bonferroni p < 0.05) displayed. Cell types most significantly correlated with CD8+ and CD4-1 cell types highlighted in yellow. (b) Circos plot showing significantly correlated (Bonferroni p < 0.05) cell types with plasma P03 and P04 cells from gastric tumors within publicly available single-cell RNA sequencing data (GSE206785). (c) Correlation of activated CD4+ T-cells with HLA-DR expression on tumor-specific cell types as determined by broad immune profiling, plotted by both strength of correlation (R) and p-value for non-zero slope. (d) Correlation of activated CD4+ T-cells with HLA-DR expression on either macrophages (above) or CD45-cells (bottom), stratified by immune subtype. Correlation (R) and p-value for non-zero slope as indicated. (e) Predictive cell type markers for response to pembrolizumab identified by regularized logistic regression. Cell type compositions inferred by CIBERSORT deconvolution of bulk sequencing data. Predictive cell types (CD4-memory/activated T-cells, M1 macrophages, CD8, CD4-memory/resting) marked at the dashed vertical line. (f) Frequency of pre-treatment CD4-memory/activated T-cells and M1 macrophages in responders and non-responders to pembrolizumab as part of NCT#02589496 (left) and frequency of equivalent immune cell populations within broad immune panel stratified by immune subtype (right). P values calculated by Wilcoxon’s rank-sum test.

Correlation analysis also noted a particularly strong association of activated CD4+ T-cells with both dendritic cells (r=0.74, Bonferonni, p=0.04) and monocytes (r=0.76, Bonferonni, p=0.04) (**Fig. 5A**), consistent with the known expression of MHC class II on these cell subsets. To explore this further, we assessed whether the presence of activated T-cells correlated with HLA-DR expression on individual cell types within the gastric cancer microenvironment. Indeed, we found that activated CD4+ T-cell frequency correlated most strongly with HLA-DR expression on macrophages and CD45-cells, and that this correlation was highest in tumors with higher IS scores (**Fig. 5C-D and Supplementary Fig. 5B**). Deep T-cell immune phenotyping showed that tumor-specific HLA-DR expression correlated most highly with proliferating and exhausted CD4+ T-cell subsets, but also with activated Tregs (**Supplementary Fig. 5C**).

Finally, to establish the clinical relevance of these newly described immune subtypes (IS), we analyzed pre-treatment bulk RNA-sequencing data from patients with metastatic gastric cancer who subsequently received single-agent pembrolizumab as part of a clinical trial (NCT#02589496).^29^ We estimated cell-type abundance from bulk RNA-seq data using CIBERSORT (**Figure 5E**).^30, 31^ Using a regularized log-ratio logistic regression model^32^ to determine which cell types were most predictive of response to pembrolizumab, we identified “CD4 memory-activated” and “M1 Macrophage” cell types as most predictive of clinical response, while “CD4 resting” cells were associated with lack of treatment response (**Fig. 5E-F**). Comparative analysis of these cell types with our broad immune panel indicated that CD4-activated, CD4-1, macrophage and monocyte clusters from our spectral cytometry data best corresponded to these inferred cell types from the CIBERSORT analysis. Interestingly, abundance of these CD4 and monocyte/macrophage cell clusters increased progressively from least to most immunogenic subtype (**Fig. 5F**). This similar pattern indicates a potential association between our newly developed immune score, a CD4+ T-cell-macrophage network, and clinical response to immune checkpoint blockade.

## Discussion

The efficacy of immune-based strategies for solid tumors has been thought to be restricted to highly immunogenic tumors based on either high somatic mutational burden or evidence of oncogenic viral transformation. Even in these tumor types, the ability of established biomarkers, including PD-L1 expression by immunohistochemistry, mutational burden, and/or tumor molecular subtype has been limited. Gastric cancer is notable for a high rate of tumor-specific PD-L1 expression,^33–36^ and immune-based strategies such as anti-PD-1 therapy have shown clinical promise,^29, 37, 38^ but identifying patients with potentially immunogenic tumors has been limited by a lack of understanding of the gastric cancer-specific immune landscape.

Our study demonstrates that current GC classification systems are insufficient in adequately identifying tumors with robust immune signatures. Our comprehensive analysis of GC immune profiles on multiparametric immune phenotyping identified three classifications of GC. These immune subtypes offer insights into the unique immune landscape of gastric cancer. First, we found that more immunogenic tumors were characterized by high rates of macrophages and plasmablasts. Several studies have shown that both cell types can initiate and sustain T-cell priming within the tumor microenvironment, and strategies to enhance recruitment of these cell types may facilitate T-cell priming in less immunogenic gastric cancer subtypes.^39–42^ This is particularly notable given that elevated levels of plasmablasts have been observed in patients with non-progressing metastatic cancers, including in melanoma, lung adenocarcinoma and renal cell carcinoma,^43^ and multiple single-cell transcriptomic studies have confirmed a significant presence of B-cells with plasma cell features within the gastric tumor microenvironment.^21, 44, 45^ Our findings showing high levels of plasma cells in immunogenic groups and the significant correlation plasma cells have with CD4 and CD8 cells demonstrates that plasma cells may indeed be a marker of immunogenicity both in gastric cancer and across cancers.

Second, we identified distinct patterns of T-cell infiltration within the gastric cancer microenvironment. CD8+ T-cells were recruited to the tumor microenvironment, but less potently than in melanomas and without strong upregulation of canonical markers of T-cell exhaustion. Conversely, CD4+ T-cell recruitment and effector differentiation was robust in gastric cancers. Furthermore, we identified strong correlations between intratumoral plasmablast and CD8+ T-cell accumulation, while CD4+ T-cell accumulation correlated with MHC-II expression on macrophages and non-immune cells. Our finding that the combination of activated CD4+ T-cells and macrophages was indicative of immunogenicity in our analysis and predictive of response to immunotherapy suggests that strategies to bolster MHC-II expression and/or CD4+ T-cell recruitment may be uniquely beneficial in gastric cancer. These findings also provide a potential explanation for why PD-1 directed therapies have only limited success in this class of tumors.

A significant issue in most solid tumors is that immunotherapy, while a promising therapeutic approach both in early^46^ and late stages of disease,^47^ has had varying results among different cancer subtypes and does not have adequate biomarkers for selecting patients most likely to respond to treatment. In our study, the immune clusters we identified through spectral cytometry-based immune phenotyping did not correlate with the current prespecified groups of immunogenic and non-immunogenic GC based on TCGA classifications, or PD-L1 and TMB scores. Furthermore, traditionally non-immunogenic GC, including intestinal, mixed, and diffuse subtypes, and tumors with PD-L1 combined positive scores (CPS) <1 were present across immune clusters including the most immunogenic subtype (IS3) which had high infiltration of activated CD8+ and CD4+ T cells. Our findings indicate that the current biomarkers and classification systems for the selection of tumors that undergo immunotherapy may be excluding tumors, such as the traditionally non-immunogenic tumors which are shown in our study to have robust immune signatures, that could have the potential to respond to therapy.

In conclusion, our data greatly contributes to a better understanding of the immune phenotypes of gastric cancer. Our findings showcasing robust immune signatures in traditionally non-immunogenic tumors highlight a critical shortcoming in the current GC selection criteria for immunotherapy which may be excluding tumors with the potential to respond to therapy. Further investigation into short- and long-term outcomes based on immune signatures are necessary to confirm the clinical implications of our study.

## Acknowledgments

This research was supported in part by the institution’s Cancer Center Support Grant (P30 CA008748) from the NIH, and the Cycle for Survival-Equinox Grant (MA, TF, VES, SAV). Figures 1A and 5E were made using images from Biorender.

## Author Contributions

MA, VES, and SAV conceived the project and designed all experiments. MA, SS, and HD performed all sample acquisition. YHL and EC performed all tissue processing and spectral cytometry assays. MA and TF performed all computational and statistical analyses. LT performed all pathologic tissue review, immunohistochemical staining and analysis. MAS, VES and SAV wrote the manuscript; all authors reviewed the manuscript.

## METHODS

After Institutional Review Board approval (IRB MED-ED-20-003), fresh gastric tumor, adjacent normal, and lymph node tissue samples were collected from patients with gastric adenocarcinoma at Memorial Sloan Kettering Cancer Center (MSK). All patients were undergoing gastric resection or upper esophagogastroduodenoscopy (EGD) and diagnostic laparoscopy entailing tissue removal as part of treatment or diagnosis. Gastric resections were completed through open or robotic-assisted procedures. Biopsies of the primary gastric tumor were performed using cold endoscopic forceps. The final pathology report, including pathologic TNM stage and PD-L1 scores, were reported by a dedicated gastrointestinal pathologist (LHT). Pathologic TNM staging was reported in accordance with the 8^th^ edition of the AJCC staging system. The final pathology of all samples had confirmed gastric adenocarcinoma on the date of tissue acquisition. Full clinical information for patients were collected in a prospectively maintained clinical database.

### Sample Collection and Processing

Fresh primary tumor, adjacent normal, or lymph node tissue samples weighing 0.1–4.5 g were transferred from the operating room (OR) in phosphate buffered saline (PBS) solution and dissected into ∼1 mm^3^ pieces in PBS. Samples were digested using a GentleMACS Dissociator in 4.7 mL of R-10 solution consisting of 10% fetal bovine serum (FBS), 10 mM HEPES, 1% penicillin-streptomycin, 1% L-glutamine, and Miltenyi enzyme (Miltenyi Biotec tumor dissociation kit, human) at 37°C for 30 minutes. Tissue suspensions were filtered through a 100-μm cell strainer and washed with cold R-10 solution. Tissue suspensions were then gently centrifuged at 500 x *g* for 5 minutes. Cell pellets were resuspended in 1 mL of lysing buffer ammonium-chloride-potassium (ACK) and incubated on ice for 10 minutes, centrifuged at 500 x *g* for 3 minutes, and washed in 15 mL cold R10 solution. Cells were treated with 40 μg/mL DNase in 60 μL + 440 μL PBS + 10 μL 1M MgCl_2_ and centrifuged at 500 x *g* for 3 minutes. Pellets were resuspended in viability dye for staining.

Cell suspensions were divided equally for staining with broad immune and T cell panels. Cell suspensions were first stained using the Ghost Dye^TM^ Violet 510 viability dye to discriminate viable cells from non-viable cells. For the viability stain, 100 μL of Ghost Dye^TM^ Violet 510 viability dye (1:400 dilution in PBS) + 25 μL FcBlock (1:20 dilution in PBS) were added to the cell suspensions and pipetted to mix. Cell suspensions were then incubated at room temperature in the dark for 10 minutes. Then, 1ml of 2%FCS/PBS was added to the cell suspensions and spun down (1500rpm, 5min, RT). Supernatant was aspirated to dry pellet.

Subsequently, single cell suspensions were stained using surface markers at room temperature for 45 minutes (**Supplementary Table 2**). 25 μL of surface panel mixed in FACS/BV Buffer was added to the cell suspensions. Cell suspensions were incubated for 45 minutes at room temperature in the dark. 1ml of 2% FCS/PBS was added to the cell suspensions and spun down (1500rpm, 5min, RT). Supernatant was aspirated to dry pellet. Cells were then fixed and permeabilized using FoxP3/Transcription Factor Fixation/Permeabilization buffer (eBioscience) for 30 minutes at room temperature in the dark. 1mL of permeabilization buffer (1x in H2O) as added to the cells and spun down (1800rpm, 5min, RT). Supernatant was aspirated to dry pellet.

Cell suspensions were then stained for intracellular proteins. 50 μL of intracellular antibodies prepared in Permeabilization Buffer were added to the cells and incubated at 4°C for 16 hours in the dark room. Then 1mL of Permeabilization Buffer was added to the cells and spun down (1800rpm, 5min, RT). Supernatant was aspirated to dry pellet. Cells were then fixed using 50 μL of with 4% paraformaldehyde (PFA) in PBS for each cell suspension and incubated for 10 minutes at room temperature in the dark. Then 250 μL of FACS buffer was added to each cell suspension and samples were acquired using a Cytek® Aurora. Data was extracted using FlowJo Software into csv files.

### Mini-batch K-means cell typing

Mini-batch K-means clustering implemented by R package mbkmeans^16^ was conducted to determine cell types separately for the broad immune and T-cell specific panels. The clustering was performed for the number of cell types k = 5, 6, …, 20, with 10 random initializations for each choice of k. The final number of cell types was determined by the Elbow method on the within-cluster sum of square curve.^15^ All gated cells were included in cell clustering without downsampling. Uniform manifold approximation and projection (UMAP)^48^ was performed using R package Seurat^49^ to visualize cell type distributions in a two-dimensional space after downsampling of 10,000 cells for each tissue sample. The default settings of 30 nearest neighbors, 0.01 minimum distance, 2 components, and Euclidean distance metric, were used for UMAP visualization. For the broad immune phenotyping panel, the following markers were used in K-means cell typing and UMAP: Clec10a, CD11b, CD15, CD8, CD19, CD16, CD20, CD45, CD274, HLA-DR, XCR1, CD56, CD11c, CD14, CD88, CD3. For the T-cell panel, the following markers were used in K-means cell typing and UMAP: PD-1, CD39, TCF-1, Ki-67, CXCR5, CCR7, CD45RA, CD8, CD38, CD27, LAG-3, CD45, CD95, CD4, HLA-DR, TOX, FoxP3, T-bet, Eomes. For CD45 myeloid cells, the following markers were used in K-means cell typing and UMAP: Clec10a, CD11b, CD15, CD8, CD19, CD16, CD20, CD45, CD274, HLA-DR, XCR1, CD56, CD11c, CD14, CD88, CD3.

### Hierarchical clustering of samples based on cell composition

Sample-specific cell composition vectors were derived from cell type identities obtained by mini-batch K-means clustering. The cell composition vectors for tumor tissues were further clustered by Euclidian distance-based hierarchical clustering using R function hclust to define immune subtypes. Hierarchical clustering was separately performed for the broad immune panel, the T-cell panel, myeloid panel, and the joint broad immune and T-cell panels. The cell compositions for different immune subtypes were visualized by stacked bar charts for tumor, normal and lymph node tissue samples.

### Data analysis for derived tumoral-immunological patient subgroups

Cell type compositions in tumor, normal, and lymph node samples were compared across tumoral-immunological patient subgroups using Wilcoxon rank sum test. Summary statistics of clinical variables, including molecular gastric cancer subtypes, T stage, N stage, M stage, neoadjuvant chemotherapy, tumor regression grade (TRG), and PD-L1 categories, were calculated and visualized as stacked bar charts across tumoral-immunological patient subgroups identified from flow cytometry data using R package ggplot2.^50^ Kruskal-Wallis rank sum test or Fisher’s exact test was used to test the association between clinical variables and tumoral-immunological patient subgroups using R package gtsummary.^51^ Alluvial plots were generated by R package ggalluvial^52^ to visualize the association between patient subgroups defined by different choices of panels and molecular gastric cancer subtypes. For tumor, lymph node, and normal tissues, Pearson correlations between cell types were calculated based on log-ratio transformed cell type composition using non-CD45 cells (broad immune panel) and non-CD4/CD8 cells (T-cell panel) as the reference composition for the two panels, respectively. Two-sided correlation tests implemented by R function “cor.test” were performed for all pairwise cell type correlations against the null hypothesis of zero correlation. Bonferroni procedure^52^ was applied to adjust for multiple correlation testing, such that cell type correlations with adjusted p-value < 0.05 were regarded as significant. Significant cell type correlations were visualized by and printed on heatmaps.

### Acquisition and analysis of publicly available single-cell transcriptomics dataset GSE206785

The single-cell RNA-sequencing dataset GSE206785 was accessed from the NCBI Gene Expression Omnibus database.^21^ This publicly available dataset contains single cells from tumors and matched normal tissues of 24 treatment-naïve patients with gastric adenocarcinoma. Cell type and subtype annotations for all 111,140 sequenced single cells from tumor samples were downloaded from the NCBI database.

There were 12 annotated cell types, namely B, CD4+ T, CD8+ T, endothelial, epithelial, fibroblast, glial, innate lymphoid, mast, mural, myeloid, and plasma. We merged the compositions of endothelial, epithelial, mural, glial, and fibroblast as non-immune cells, and kept the compositions of the rest immune cell types. Pearson correlations between cell types were then calculated based on log-ratio transformed cell subtype composition using non-immune cells as reference. Two-sided cell subtype correlation tests were performed against the null hypothesis of zero correlation followed by Bonferroni correction. Cell subtype correlations with adjusted p-value < 0.05 were regarded as significant. Significant cell subtype correlations were displayed by circos plots generated by R package circlize.^53^

## Cell type imputation and predictive modeling of bulk transcriptomic data PRJEB25780

The preprocessed bulk RNA-sequencing dataset PRJEB25780 was retrieved from the Tumor Immune Dysfunction and Exclusion (TIDE) repository (https://tide.dfci.harvard.edu). This publicly available dataset consists of gene expressions for 61 patients treated with metastatic gastric cancer who received pembrolizumab as part of clinical trial NCT#02589496.^53^ We performed cell type imputation for the bulk sequencing data PRJEB25780 using CIBERSORTx online platform (https://cibersortx.stanford.edu).30 Patients without available response outcomes were excluded from the analyses. The default LM22 signature matrix file was used with B-mode batch correction, disabled quantile normalization, relative run mode, and 500 permutation runs. All but one imputed sample passed the CIBERSORTx built-in test against the null hypothesis that no cell types in the LM22 matrix were present in the bulk data. To standardize cell types defined by CIBERSORTx’s LM22 signature matrix and by our flow cytometry experiments, we applied the correspondences displayed in the following table. To account for the limitation that no non-immune cells were imputed by CIBERSORTx, we fitted a regularized logistic regression model for log-ratios between standardized imputed cell type fractions.^32^ Cell type features which obtained the lowest 5-fold cross-validated mean-squared error were selected for further comparison between response outcomes and the immune clusters defined by the flow cytometry data.

| **CIBERSORTx LM12 cell types** | **Spectral cytometry cell types (panels)** |
| --- | --- |
| B cells naive | B cells (broad immune) |
| B cells memory + Plasma cells | Plasmablast (broad immune) |
| T-cells CD8 | CD8 (broad immune) + CD8-activated + CD8-effector + CD8-eomes (T-cell) |
| T-cells CD4 naive | Not applicable |
| T-cells CD4 memory resting | CD4-2 (broad immune), CD4-memory (T-cell) |
| T-cells CD4 memory activated | CD4-1 + CD4-activated (broad immune), CD4-activated + CD4-effector + CD4-exhausted + CD4-proliferating (T-cell) |
| T-cells regulatory | T-regs (T-cell) |
| T-cells gamma delta | Not applicable |
| NK cells resting | NK cells (broad immune) |
| NK cells activated | Not applicable |
| Monocytes | m-MDSC (broad immune) |
| Macrophages M0 | Not applicable |
| Macrophages M1 | Macrophages + Monocytes (broad immune) |
| Macrophages M2 | Not applicable |
| Dendritic cells resting + Dendritic cells activated | DC (broad immune) |
| Mast cells, Mast cells activated, Eosinophils | Not applicable |
| Neutrophils | g-MDSC (broad immune) |

---

**Supplementary Table 1.**
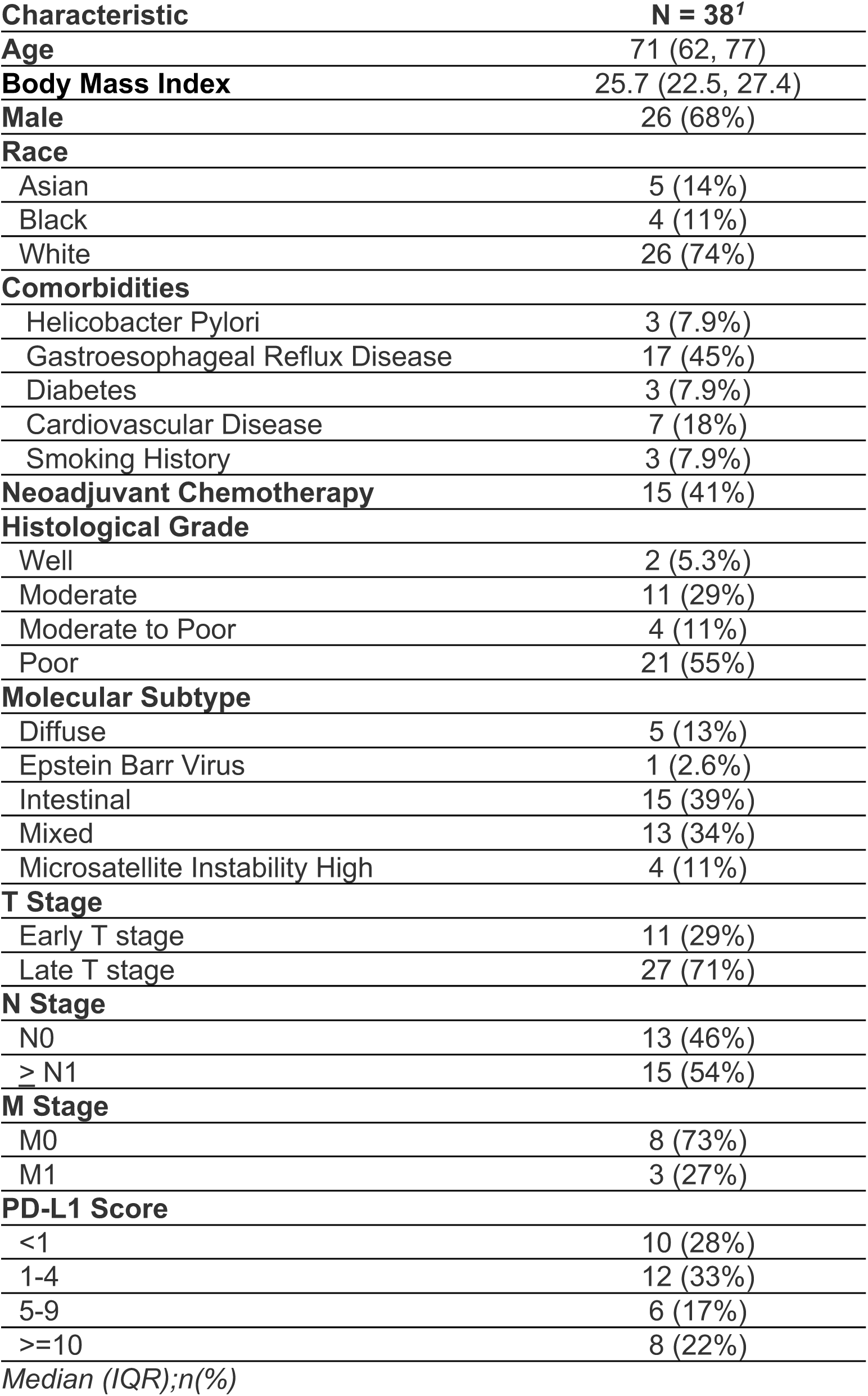
Demographics of patients with gastric cancer from whom tissue samples underwent spectral cytometry-based immune phenotyping

**Supplementary Table 2.** Antibodies used in broad immune and T-cell panels for spectral cytometry.

| Broad Immune |  |  |  | T-cell |  |  |  |
| --- | --- | --- | --- | --- | --- | --- | --- |
| Target | Clone | Fluorophore | Dilution | Target | Clone | Fluorophore | Dilution |
| CD45 | HI30 | BV570 | 1:200 | CD45 | HI30 | BV570 | 1:200 |
| CD19 | SJ25C1 | BUV563 | 1:100 | CD4 | RPA-T4 | BV711 | 1:200 |
| CD20 | 2H7 | BUV805 | 1:400 | CD8 | RPA-T8 | BUV563 | 1:200 |
| CD3 | SK7 | PerCP-Cy5.5 | 1:25 | CD45RA | HI100 | BUV395 | 1:400 |
| CD8 | RPA-T8 | BUV496 | 1:200 | CD27 | L128 | BUV737 | 1:50 |
| CD56 | 5.1H11 | PE | 1:50 | CCR7 | 3D12 | BB700 | 1:25 |
| HLA-DR | G46-6 | BV786 | 1:50 | CD95 | DX2 | BV650 | 1:50 |
| CD11c | B-ly6 | PE-Cy5 | 1:50 | CD38 | HIT2 | BUV661 | 1:200 |
| XCR1 | S15046E | FITC | 1:50 | HLA-DR | G46-6 | BV786 | 1:50 |
| Clec10a | H037G3 | APC | 1:200 | CD39 | A1 | APC-Fire 750 | 1:50 |
| CD14 | 63D3 | PE-Cy7 | 1:50 | PD-1 | EH12.2H7 | APC | 1:100 |
| CD11b | LM2 | Alexa 647 | 1:50 | LAG-3 | 11C3C65 | BV421 | 1:100 |
| CD88 | S5/1 | PE-Dazzle594 | 1:100 | Ki-67 | B56 | Alexa 700 | 1:200 |
| CD274 | 29E.2A3 | BV650 | 1:100 | FoxP3 | PCH101 | PE-Cy5 | 1:100 |
| CD16 | 3G8 | BUV615 | 1:100 | CXCR5 | RF8B2 | BB515 | 1:25 |
| CD15 | HI98 | Alexa 700 | 1:50 | TCF-1 | C63D9 | Alexa 488 | 1:100 |
|  |  |  |  | TOX | REA473 | PE | 1:50 |
|  |  |  |  | T-bet | 4B10 | PE-Cy7 | 1:400 |
|  |  |  |  | Eomes | WD1928 | PE-eFluor 610 | 1:100 |

