## Supplementary Figures for "High-Dimensional Spectral Cytometry Reveals Therapeutically Relevant Immune Subtypes in Gastric Cancer"

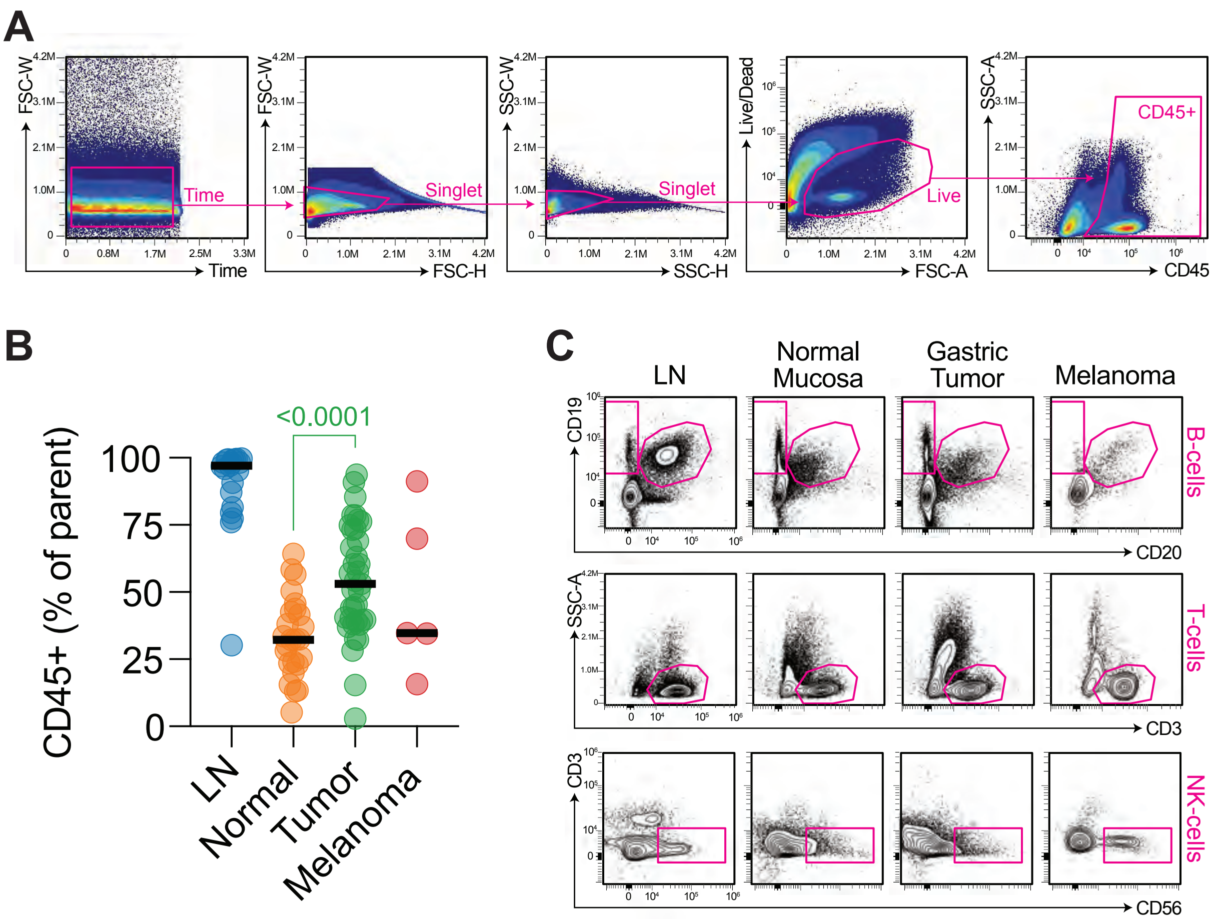


**Supplementary Figure 1.** Broad immune phenotyping of gastric adenocarcinoma. (a) Gating strategy for broad immune panel. (b) Proportion of CD45+ immune cells in gastric tumors, normal mucosa, lymph nodes, and melanoma tumors. (c) Contour plots of lymphocyte cell subtype abundances in gastric tumors, normal mucosa, and lymph nodes compared with melanoma tumors. P values as indicated for gastric tumors were calculated by one-way ANOVA with Sidak's multiple comparisons post-test relative to normal mucosa.

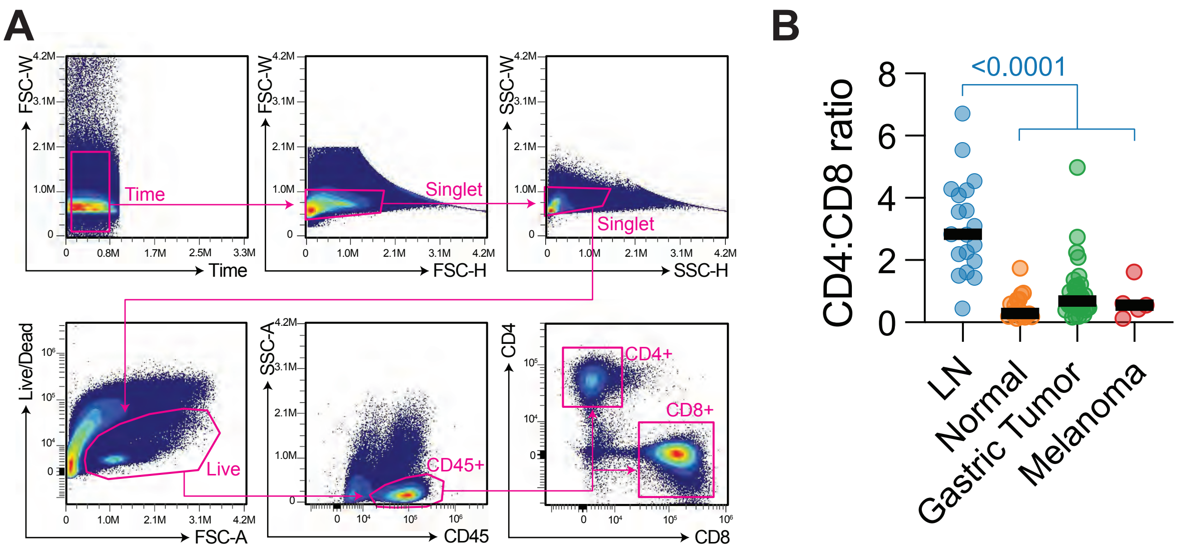


**Supplementary Figure 2.** T-cell immune phenotyping of gastric adenocarcinoma. (a) Gating strategy for T-cell immune panel. (b) Ratio of CD4:CD8 T-cells in gastric tumors, normal mucosa, lymph nodes, and melanoma tumors. P values as indicated for lymph nodes were calculated by one-way ANOVA with Sidak's multiple comparisons post-test relative to normal mucosa, gastric tumors, and melanoma tumors.

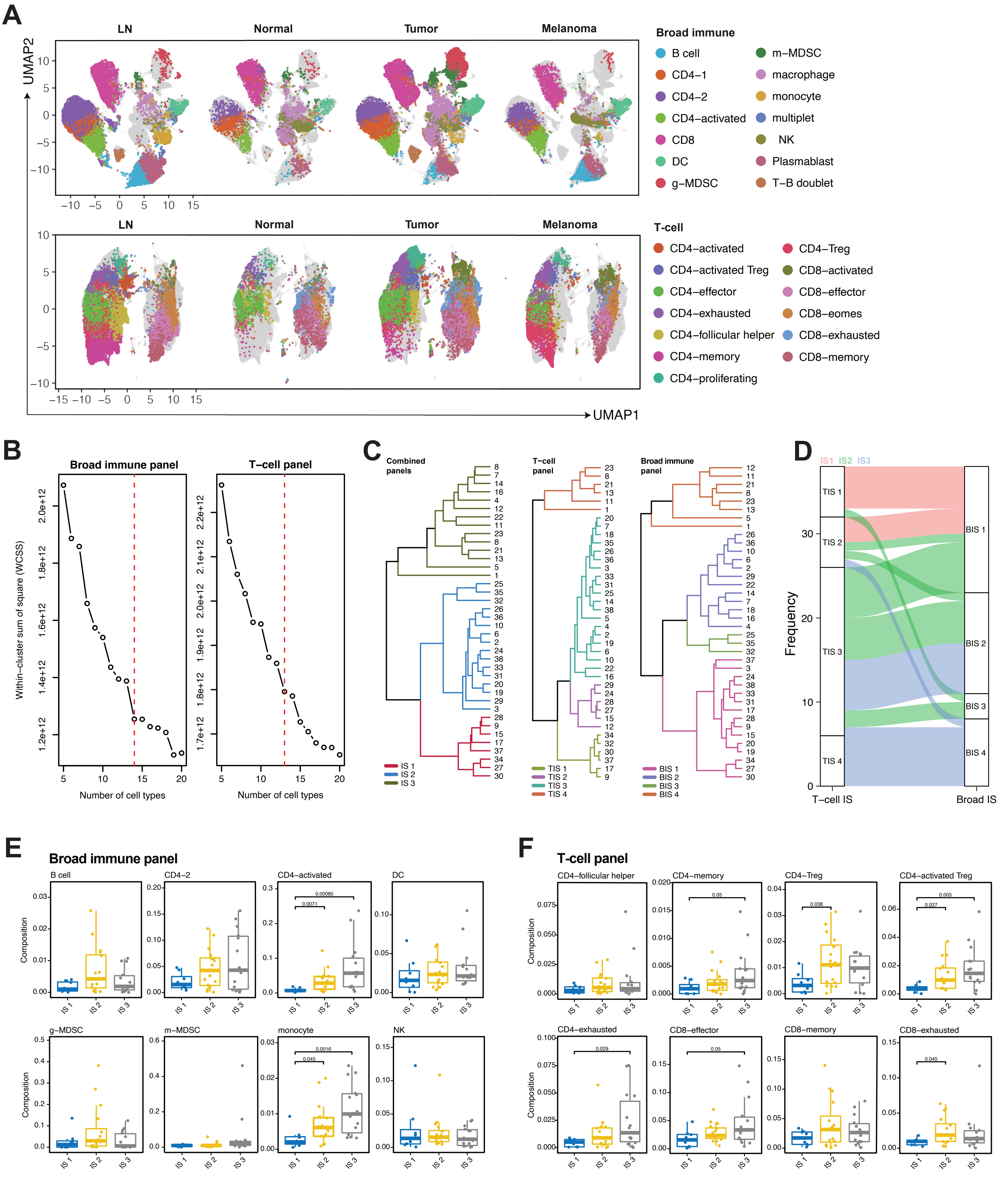
A
LN
Normal
Tumor
Melanoma
UMAP2
UMAP1
Broad immune
B cell
CD4−1
CD4−2
CD4−activated
CD8
DC
g−MDSC
m−MDSC
macrophage
monocyte
multiplet
NK
Plasmablast
T−B doublet
T-cell
CD4−activated
CD4−activated Treg
CD4−effector
CD4−exhausted
CD4−follicular helper
CD4−memory
CD4−proliferating
CD4−Treg
CD8−activated
CD8−effector
CD8−eomes
CD8−exhausted
CD8−memory
B
Broad immune panel
T−cell panel
Within−cluster sum of square (WCSS)
Number of cell types
C
Combined panels
T−cell panel
Broad immune panel
IS 1
IS 2
IS 3
TIS 1
TIS 2
TIS 3
TIS 4
BIS 1
BIS 2
BIS 3
BIS 4
D
IS1 IS2 IS3
Frequency
T−cell IS
Broad IS
E
Broad immune panel
B cell
CD4−2
CD4−activated
DC
g−MDSC
m−MDSC
monocyte
NK
Composition
0.00085
0.0071
0.0016
0.045
F
T-cell panel
CD4−follicular helper
CD4−memory
CD4−Treg
CD4−activated Treg
CD4−exhausted
CD8−effector
CD8−memory
CD8−exhausted
Composition
0.05
0.038
0.005
0.027
0.029
0.05
0.045

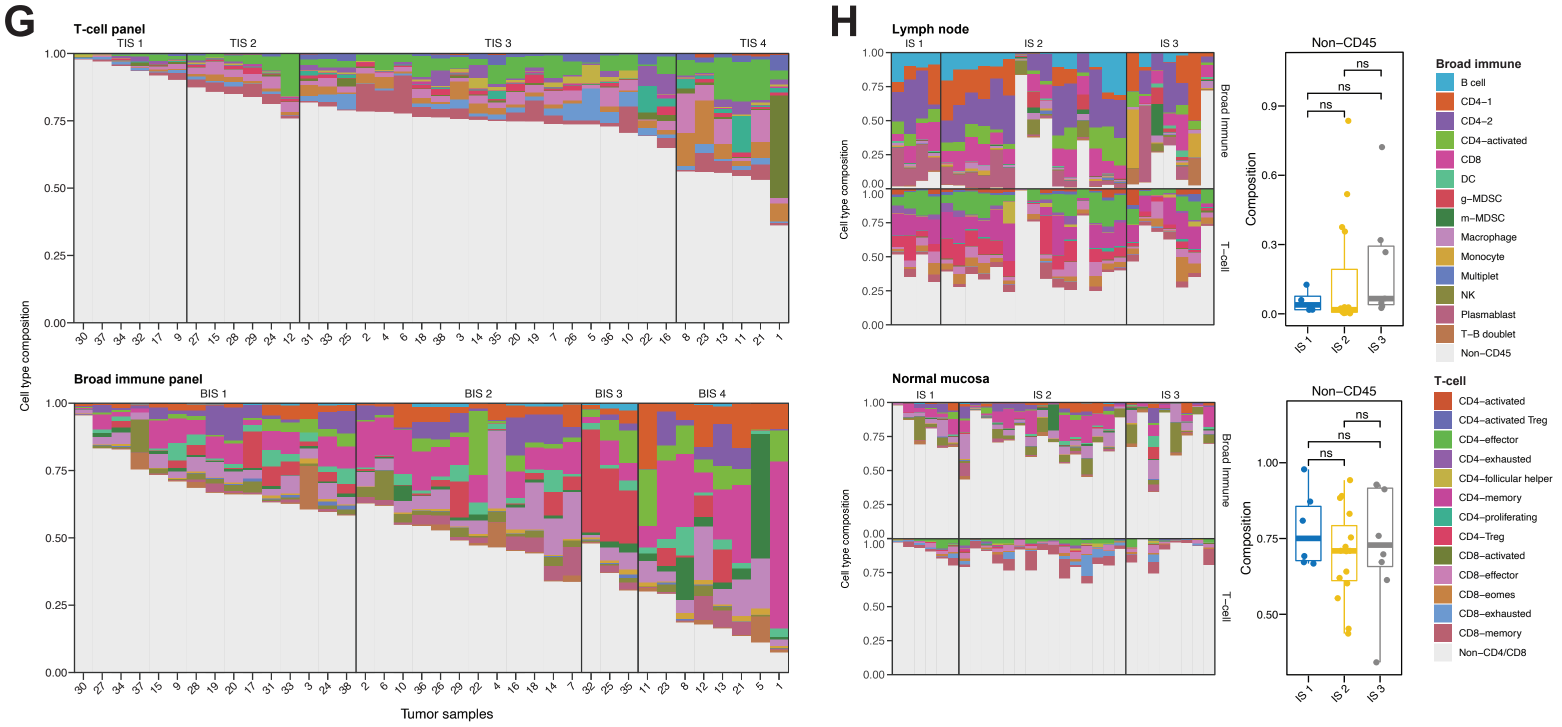


**Supplementary Figure 3.** Construction and composition of immune subtypes in gastric adenocarcinoma. (a) UMAP projection of CD45+ immune cells based on broad immune panel (above) and CD4+ and CD8+ lymphocytes based on T-cell panel (below), stratified by tissue type. Cell type annotations as indicated. Grey indicates UMAP project of cells integrated across tissue types. (b) Within-cluster sum of square of mini-batch K-means clusters versus number of clusters for broad immune panel (left) and T-cell panel (right). Selected number of cell types denoted by red dashed vertical line by the elbow method. (c) Hierarchical clustering dendrograms for combined broad immune and T-cell panels (left), T-cell panel (middle) and broad immune panel (right). (d) Alluvial plots showing concordance between T-cell and broad immune subtypes. (e,f) Frequency of cell type by immune score as a proportion of viable cells using broad immune panel (e) or T-cell panel (f). (g) Distribution of patient-specific cell type frequency within tumors stratified by T-cell specific immune subtype (above) or broad immune phenotype (below). (h) Left, distribution of patient-specific cell type frequency within lymph nodes (above) or normal mucosa (below), stratified by immune subtype. Right, frequency of nonimmune cells as a proportion of viable cells in lymph nodes (above) and normal mucosa (below), stratified by immune score. P values as indicated were calculated by Wilcoxon's rank-sum test.

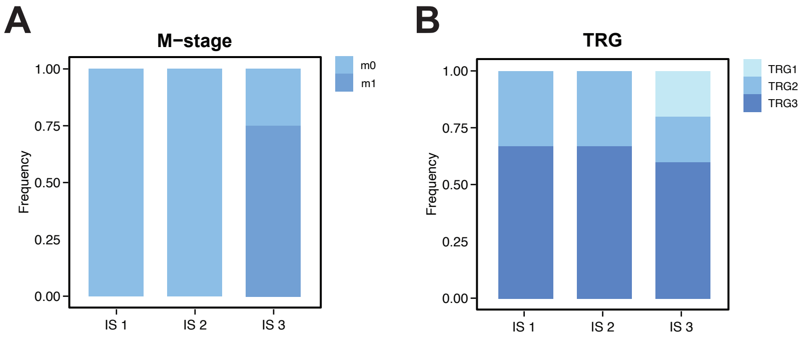


**Supplementary Figure 4.** Gastric cancer immune subtypes are independent of pathologic stage and response to Neoadjuvant Chemotherapy. (a) Distribution of pathologic M stage across immune subtypes. (b) Distribution of tumor response grade (TRG) across immune subtypes. TRG1 = 100% response, TRG2 = less than 10% residual tumor, TRG3 = 10-50% residual tumor.

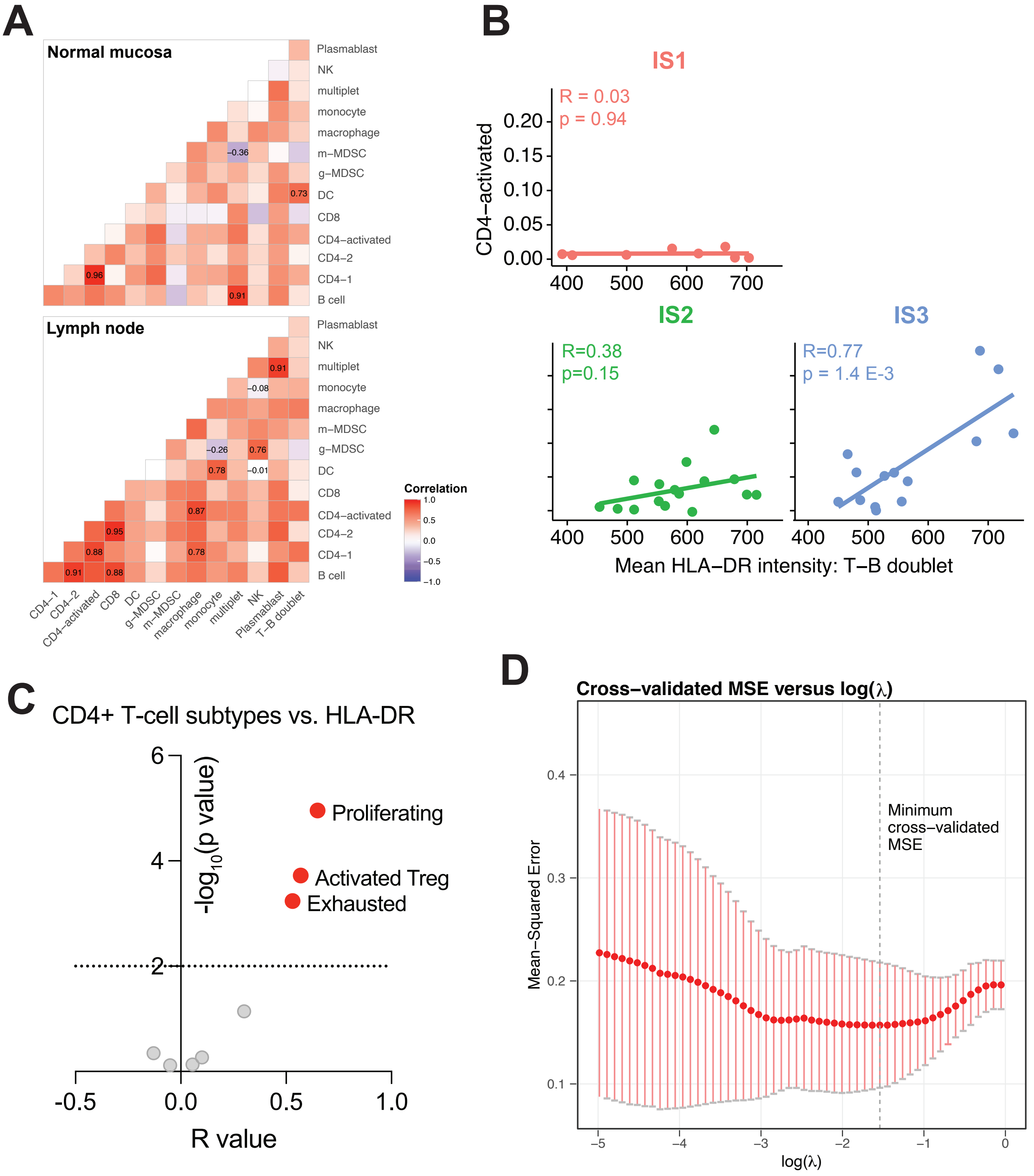


**Supplementary Figure 5.**

**Supplementary Figure 5.** Immunogenic gastric tumors are marked by tumor-specific CD4+ T-cell activation. (a) Correlation analysis of immune cell types within normal mucosa (above) and lymph nodes (below) based on broad immune phenotyping panel. Correlation values significantly different from zero (Bonferroni $p < 0.05$) displayed. (b) Correlation of activated CD4+ T-cells with HLA-DR expression on T-cell/B-cell conjugates, stratified by immune subtype. Correlation (R) and p-value for non-zero slope as indicated. (c) Correlation of CD4+ T-cell subtypes based with HLA-DR expression across viable non-CD4 cells measured using the T-cell immune profiling panel, plotted by both strength of correlation (R) and p-value for non-zero slope. (d) Cross-validated mean square error (MSE) versus choices of penalty parameters by regularized logistic regression for response to pembrolizumab. Penalty parameter with smallest cross-validated MSE annotated by dashed vertical line.
